# Salivary exosomal miR-1307-5p predicts disease aggressiveness and poor prognosis in oral squamous cell carcinoma patients

**DOI:** 10.1101/2022.07.13.499918

**Authors:** Aditi Patel, Shanaya Patel, Parina Patel, Dushyant Mandlik, Kaustubh Patel, Vivek Tanavde

## Abstract

**Background:** Salivary exosomal miRNAs as biomarkers facilitate repeated sampling, real-time disease monitoring and assessment of therapeutic response. This study identifies a single salivary exosomal miRNA prognosticator that will aid in improved patient outcome using a liquid biopsy approach.

**Method:** Small RNA and transcriptome sequencing profiles of tumour tissues and salivary exosomes from oral cancer patients were compared to their non-cancerous counterparts. We validated these results using the Cancer Genome Atlas database and performing Real-time PCR on a larger patient cohort. Potential target genes, miRNA-mRNA networks and enriched biological pathways regulated by this microRNA were identified using computational tools.

**Results:** Salivary exosomes (size: 30-50nm) demonstrated a strong expression of CD47 and detectable expression of tetraspanins CD63, CD81 and CD9 by flow cytometry. miR-1307-5p was exclusively overexpressed in tissues and salivary exosomes of oral cancer patients compared to their non-cancerous counterparts. Enhanced expression of miR-1307-5p clinically correlated with poor patient survival, disease progression, aggressiveness and chemo-resistance in these patients. Transcriptome analysis suggested that miRNA-1307-5p could promote oral cancer progression by suppressing *THOP1, EHF, RNF4, GET4*, and *RNF114*.

**Conclusion:** Salivary exosomal miRNA-1307-5p is a potential prognosticator for predicting poor survival and poor patient outcome in oral cancers.

## Introduction

Oral cancer is one of the most commonly occurring sub-type of head and neck cancers with high mortality and recurrence rates. Asia accounts for 48.7% of the total cases of oral squamous cell carcinoma (OSCC) reported globally (1). In spite of recent advances in treatment strategies, factors like metastasis, loco-regional invasion, and therapeutic refractoriness significantly influence the poor prognosis of patients and ultimately contribute to a low 5-year overall survival rate. Determining the loco-regional aggression and the status of metastasis during the initial staging of the tumour may improve the overall survival and reduce the administration of aggressive treatments in the later course of the disease. The conventional approach for determining the aggressiveness of the tumour is completely reliant on imaging technologies. The rates of regional, locoregional, and local recurrences in OSCC patients are 31.2–62.6%, 4.1–16.3%, and 24–51.1% respectively which ultimately require patients to undergo repetitive diagnostic processes (2). Thus it is necessary to understand the key factors driving OSCC prognosis and identify robust markers that may act as prognosticators for the disease.

Tumours shed exosomes and provide dynamic information about the tumour mass at the sample collection time point (3). Emerging evidence suggests that tumour-derived exosomes are essential molecules that promote the formation of a pre-metastatic niche by transporting activators of epithelial-mesenchymal transition (EMT) (miRNAs, mRNAs, lncRNAs, etc.) to distant sites (4, 5). Circulating exosomal miRNAs (exomiRs) are packed into extracellular vesicles and are not prone to RNase-mediated degradation. Hence they are more stable in various biofluids, like saliva, and are tissue-specific, thereby making them attractive biomarkers for cancer prognosis. Liquid biopsy is a non-invasive biopsy method which may facilitate repeated sampling, and real-time monitoring of OSCC while allowing clinicians to observe the therapeutic response of patients. Saliva is the ideal biofluid for liquid biopsies in OSCC patients as it exists in close proximity to the tumour. The potential of exomiRs as markers for disease progression, aggression, and risk prediction has been explored across various malignancies (6). Previous studies have identified certain salivary exomiRs like miR-486-5p, miR-134, miR-24-3p, and miR-200a that can be used as markers for early detection of oral cancer (7–9). However, all of these exomiRs have been identified as biomarkers for early detection or screening of various oral cancer subsites including tongue, pharynx, and hypopharynx. The utility of salivary exomiRs as markers of OSCC progression, aggression, and prognosis is still unexplored. Furthermore, these studies have solely focused on the expression profiles of the aforementioned miRNAs. However the underlying mechanisms responsible for tumour progression, governed by these miRNAs have not been explored.

The present study identifies miR-1307-5p as a novel candidate miRNA that is exclusively expressed in OSCC samples. Further, our data demonstrates that this miRNA may be a useful biomarker for predicting disease progression and prognosis. Additionally, we have proposed a mechanism via which this miRNA may regulate chemoresistance and progression of OSCC.

## Methodology

### Patient cohort and sampling details

Samples of resected tumour tissue and unstimulated saliva were collected from patients diagnosed with oral squamous cell carcinoma (OSCC) (excluding tongue, larynx, pharynx and hypopharynx) from HCG Cancer Centre, Ahmedabad. Brush biopsies and unstimulated whole saliva of healthy individuals (matched for age and gender with the patients) with no etiological history of tobacco chewing and no clinically detectable oral lesions were used as normal controls. Resected tissue specimens were processed for histopathological evaluation and additional tumour samples were snap-frozen and stored at −80°C for further processing. Whole saliva was collected into sterile tubes from healthy individuals, and OSCC patients prior to their surgical resection according to the widely-used protocol for the collection of saliva (10). We divided the participants into a discovery cohort, consisting of 4 buccal scrapings (pooled) and 3 salivary exosomes from controls (pooled), 12 OSCC tissue samples and 8 salivary exosome samples from OSCC patients. We used pooled samples for controls to generate a reference value for controls from multiple healthy volunteers. For buccal scrapings we pooled 4 patients per sample and for salivary exosomes we pooled 3 volunteers per sample. The validation cohort consisted of 5 control buccal scrapings (pooled; 3 volunteers per pooled sample), 5 salivary exosomes from healthy volunteers, 19 OSCC tissue samples and 12 OSCC saliva samples. Demographic information and clinical data of the patients have been represented in Table 1. Paediatric cases, HCV, HPC, HIV, and HbsAg positive malignancies, and patients treated with neoadjuvant and/or adjuvant therapeutic modalities were excluded from the study. Written informed consent for participation in this study has been provided by all the patients. This study was reviewed and approved by the HCG Cancer Centre’s Ethics Committee (ECR/92/Inst/GJ/2013/RR-16) for human subject research and the use of human tissues and it also complies with the guidelines set forth by the Declaration of Helsinki (2008).

**Table 1:**
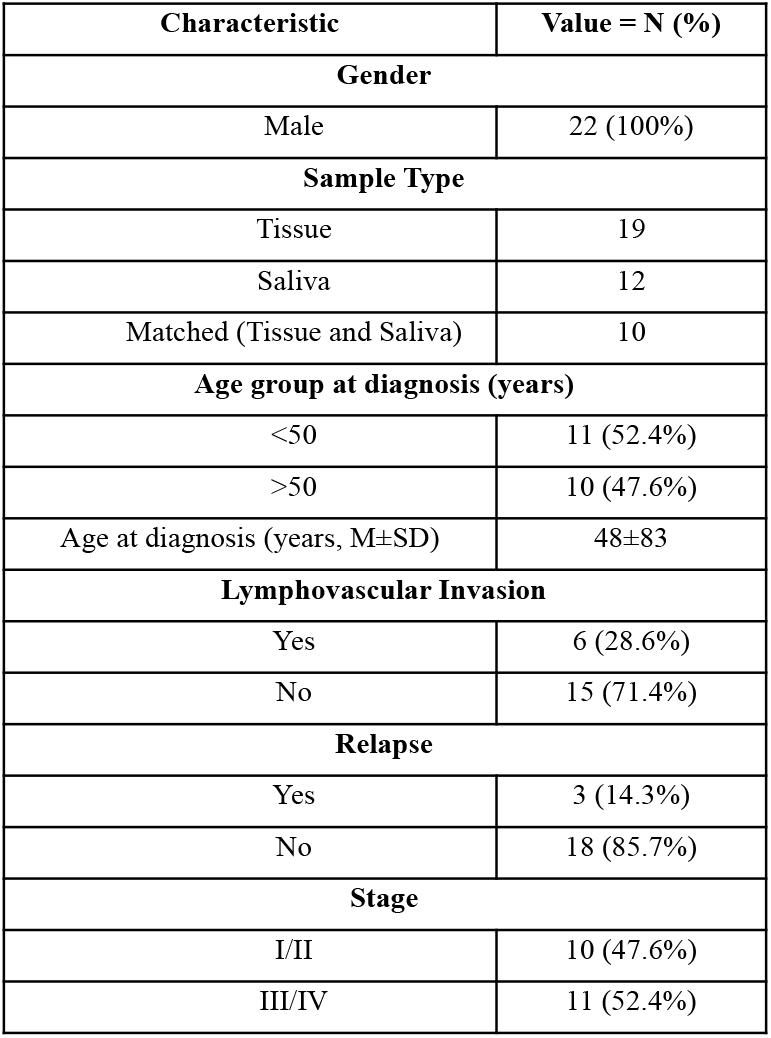
The clinicopathologic characteristics in OSCC patients

| Characteristic | Value = N (%) |
| --- | --- |
| **Gender** |  |
| Male | 22 (100%) |
| **Sample Type** |  |
| Tissue | 19 |
| Saliva | 12 |
| Matched (Tissue and Saliva) | 10 |
| **Age group at diagnosis (years)** |  |
| <50 | 11 (52.4%) |
| >50 | 10 (47.6%) |
| Age at diagnosis (years, M±SD) | 48±83 |
| **Lymphovascular Invasion** |  |
| Yes | 6 (28.6%) |
| No | 15 (71.4%) |
| **Relapse** |  |
| Yes | 3 (14.3%) |
| No | 18 (85.7%) |
| **Stage** |  |
| I/II | 10 (47.6%) |
| III/IV | 11 (52.4%) |

For validation of the findings of this study, miRNA-sequencing data from the TCGA-HNSC dataset (subsites: floor of mouth, gum, palate, and other and unspecified parts of the mouth) from NIH National Cancer Institute Genomic Data Commons Data Portal (The Cancer Genome Atlas; https://cancergenome.nih.gov/) was downloaded. miRNA sequencing results and clinicopathological and survival data of 114 oral cancer patients was included in our analysis.

### Exosome Isolation

Saliva of OSCC patients and healthy controls was diluted with PBS (1:1) followed by centrifugation at 2000g for 10 minutes at room temperature. Exosomes were precipitated from the supernatant using Invitrogen™ Total Exosome Isolation Reagent from other body fluids kit (Thermo Fisher Scientific, Waltham, MA, USA) according to the manufacturer’s protocol. The precipitated exosomes were re-suspended either in TRIzol LS reagent (Thermo Fisher Scientific, Waltham, MA, USA) or in PBS depending on the subsequent analysis to be performed.

### Exosome Characterization

The number of exosomes and their sizes were determined by conducting Nanoparticle Tracking Analysis (NTA) on NanoSight LM10 (Malvern Instruments Ltd) after diluting the exosome pellet in a physiologic solution (1:500). Imaging of negatively stained EVs (NanoVan, Nanoprobes, Yaphank, NY, USA) was carried out via transmission electron microscopy (TEM) using the Jeol JEM 1010 electron microscope (Jeol, Tokyo, Japan). For imaging, exosomes were set onto 300 mesh carbon formvar grids followed by an incubation with 5 μL 2% glutaraldehyde for 3 min. The excessive fluid was wick dried and the grids were rinsed with water. Samples were stained with 2% uranyl acetate for 1 min followed by removal of excessive dye and air-dried for 10 minutes. The samples were then examined at 100kV on the transmission electron microscope at the Centre for Excellence in Imaging, Ahmedabad University.

Isolated exosomes were also characterised by flow cytometry using the vFC EV Analysis kit (Cellarcus Biosciences, San Diego, CA) according to manufacturer’s instructions. Briefly, samples were stained with a membrane stain (vFRed, Cellarcus Biosciences, CA, USA). Incorporation of a lipophilic membrane dye like vFRed in the protocol helps us to discriminate between exosomes and electronic noise. The instrument was then calibrated using a vesicle size standard of synthetic 50 nm liposomes (Lipo50; Cellarcus Biosciences, CA, USA). Since coincidence of events is a major challenge during acquisition of exosomes, we measured the event rate during acquisition of serially diluted samples. The lowest dilution where the event rate dropped in proportion with sample dilution was considered the optimal dilution. This optimal dilution was 1:80 for our salivary exosomal samples from OSCC patients. The samples (at optimal dilution) were then incubated with PE tagged antibodies against tetraspanins (TS; CD9, CD63, and CD81) (Cellarcus Biosciences, CA, USA) and APC tagged CD47 antibody (Thermo Scientific, Waltham, MA, USA) for one hour at room temperature. Samples were acquired on the CytoFLEX-LX (Beckman Coulter, Brea, CA, USA) flow cytometer. Thresholding was set on the violet laser side scatter (v-SSC) and voltages were set as per instructions in the Cellarcus protocol. All samples were acquired using the high flow rate for 2 minutes. We used a hierarchical gating strategy to identify the exosomes expressing CD47 and tetraspanins. Since our exosomes expressed CD47 strongly, we used CD47 and vFRed as primary discriminators. Double positive events were then analysed for expression of tetraspanins and events stained with vFRed, CD47 and tetraspanins were considered as exosomes.

### RNA extraction

#### Exosomes

RNA from salivary exosomes was isolated using the method described by Prendergast et al. RNA was extracted by adding 750μL TRIzol LS reagent (Thermo Fisher Scientific) and 200μL chloroform to 40μL exosomal sample. We used 5μL glycogen (5 mg/mL, Sigma Aldrich, St. Louis, MO) instead of 3μL in the original protocol. The RNA pellet was resuspended in 32μL of nuclease-free water. The final concentration of the purified RNA was determined using Qubit Fluorometer 4.

#### Tissues

TRIzol reagent was used to extract total RNA from OSCC tissue (as per manufacturer’s protocol). Purity and yield of extracted RNA were determined using Agilent 2100 Bioanalyzer (Agilent Technologies, Santa Clara, CA, United States) and Qubit Fluorometer 4 (Thermo Fisher Scientific, Waltham, MA, United States).

### miRNA Sequencing

The extracted total RNA from the samples was submitted for miRNA sequencing via Illumina NextSeq500 platform. Small RNA library preparation and sequencing were performed by Genotypic Technology Pvt. Ltd., Bangalore, India. Small RNA libraries were prepared using the QIAseq^®^ miRNA Library Kit (Cat: 331502). The ligation of 3’ and 5’ adapters to the total RNA was followed by reverse transcription and amplification of the RNA using polymerase chain reaction (PCR) which resulted in the enrichment and barcoding of cDNA. Fragment size distribution of the libraries was followed by sequencing (12-15 million single end reads per run, per sample). FastQC was used to assess the overall quality of raw sequencing reads and Cutadapt (v. 1.8) was used for trimming the reads (11, 12). The filtered reads were mapped to the human reference genome (GRCh, version 38) using miRDeep2 (v0.0.7). The same toolkit was used to annotate potential novel miRNAs (cut-off: 6) and validate the predicted miRNAs from miRbase. DESeq2 was used to conduct statistical analysis of genes with read counts in ≥ 30 of the samples. A likelihood ratio test was conducted to contrast the conditions (leucoplakia-control)-(tumor-control) and obtain differentially expressed miRNAs among 3 levels (log2 FC>2; adjusted P <0.01).

### Transcriptome Sequencing

RNA extracted from tumour tissues and salivary exosomes of OSCC patients was subjected to mRNA-sequencing wherein library preparation and sequencing was performed by Genotypic Technology Pvt. Ltd., Bangalore, India. cDNA libraries were generated using Illumina TruSeq RNA Sample Preparation v2 Kit (Lot# RS-122-2001, RS-122-2002) followed by paired-end sequencing on the Illumina HiSeq 2500 (2×150bp). Raw sequencing reads were subjected to quality assessment using FastQC followed by subsequent cleaning via Cutadapt. The refined reads were compared with a reference genome (GRCh, version 38) using Subread aligner.

### miRNA Target Prediction

In order to identify the gene targets of miRNA, TargetScan v8.0 prediction algorithm was used. Genes were considered differentially expressed if FC ≤ 2 and confidence was measured using predictive relative KD ≤ −2.

Pearson correlation and linear regression analysis were used to determine the correlation between the expression of miRNA and its target genes. For this, the RNASeq expression data was combined with the TCGA_HNSC expression data. Pearson correlation coefficient was calculated in R and the regression plots were made using the ggplot2 library in R.

### Statistical Analysis

Receiver Operating Characteristic (ROC) curves were used to determine the discrimination power of the differentially expressed miRNAs as biomarkers for OSCC aggression and progression. GraphPad Prism 6.01 software was used for constructing the curves and the sensitivity, specificity, area under curve (AUC) and p value were calculated and the optimal threshold value was decided using Youden’s index (sensitivity+specificity −1).

Survival analysis and t-tests to determine statistical significance were performed using SPSS software (v19.0; SPSS, Inc. Chicago, IL) and GraphPad prism 9.0 respectively. The results are presented as mean ± standard deviation (SD) with a student’s t-test. All experiments were repeated thrice and P < 0.05 was the threshold to determine statistical significance.

## Results

### Identification and characterization of salivary exosomes derived from oral cancer patients

Exosomes were isolated from unstimulated saliva samples of patients diagnosed with OSCC and healthy individuals. The identification and characterization of exosomes was focused on various criteria such as size distribution and concentration using Nano-sight Tracking Analysis (NTA), validation of spherical morphology by Transmission Electron Microscopy (TEM) and the presence of exosome markers revealed by flow cytometry.

NTA suggested that salivary exosomes appeared as a homogenous population with mean size of 32.9 nm and concentration of 3.66 x 10^8^ particles/ml (Figure 1(a)). Further TEM results represented the sphere-shaped morphology of the vesicles with a size ranging from 40-50 nm which was consistent with NTA profiles (Figure 1(b)). To assess the purity of the isolated exosomal population, the presence of CD47, CD63, CD9 and CD81 markers was estimated using flow-cytometric analysis. We initially calibrated the flow cytometer using 50 nm synthetic liposomes stained with vFRed, a dye that binds to lipid membrane. Salivary exosomes showed a similar violet-SSC and vFRed staining pattern as compared to the Lipo-50 synthetic liposomes indicating a size around 50 nM. These exosomes showed a strong expression of CD47. We were also able to detect the expression of the tetraspanins CD9, CD63 and CD81 characteristic of exosomes (Figure 1(c,d,e)). Collectively, these findings substantiated that the vesicles derived from patients and healthy volunteers derived saliva constituted a pure exosomal population.

**Figure 1:**
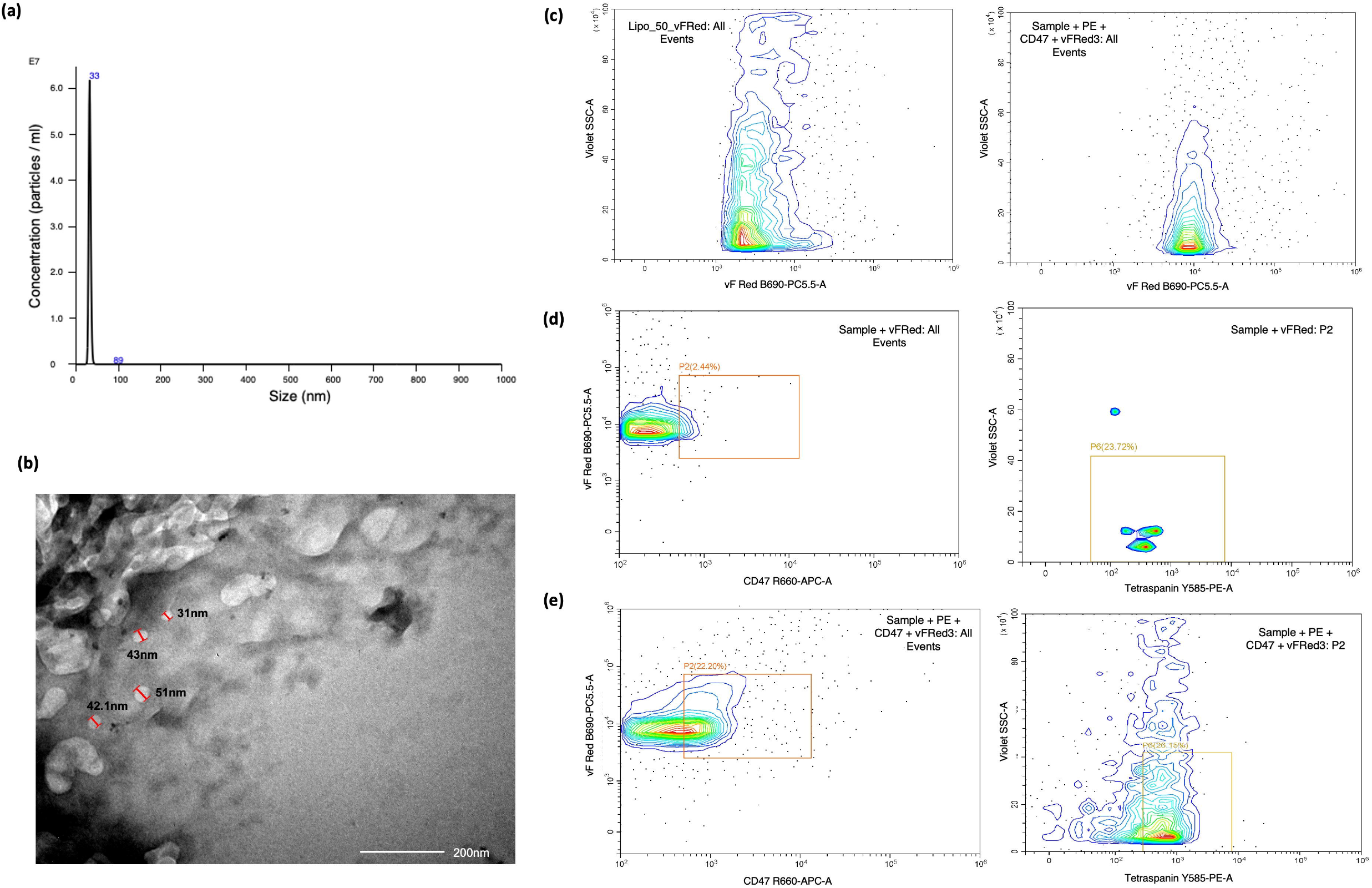
Characterization of exosomes based on size, scatter and antigen expression. (a) Representative TEM images of exosomes from OSCC patients, (Scale: 200nm); (b) Estimation of size and concentration of salivary exosomes from OSCC patients using Nanoparticle Tracking Analysis (NTA). The x-axis indicates the size distribution of particles and the y-axis shows the signal intensity in NTA; *(c-e)* Characterization of exosomes by flow cytometry. We used a membrane-binding dye vFRed to stain exosomes and examined the violet side scatter v/s vFRed staining of these exosomes. This profile was similar to the profile of Lipo-50 liposomes used as a reference standard. Salivary exosomes were stained with antibodies against tetraspanins (CD63, CD81 and CD9) labelled with PE, and APC labelled CD47 antibody. We identified a distinct CD47+/vFRed+ population in the exosome population that was absent in the unstained control. This population was then examined for tetraspanin expression and showed detectable levels of tetraspanins. Since these particles stain with vFRed, have a violet side scatter similar to 50 nm liposomes, express CD47 and tetraspanins, we concluded that these are exosomes.

### Small RNA sequencing analysis identifies miR-1307-5p, a miRNAexclusively expressed in tissues and salivary exosomes of OSCC patients

A preliminary focus of this study was to identify miRNAs exclusively expressed in tumor and salivary exosomes derived from OSCC patients with respect to their age-sex matched normal counterparts. We performed small RNA sequencing of the aforementioned discovery cohort. In OSCC patient derived tumour tissue and salivary exosomes, with a 260/280 ratio ranging from 1.87-2.12 (average 2.06) and an average RNA concentration of 655.02 ng/μl and 57.31 ng/ μl respectively. After pre-processing, adapter sequence trimming and filtering low quality reads, the mean q20 values for tissue based libraries were 97.6% (2,31,53,568) whereas salivary exosome derived libraries demonstrated a mean q20 value of 93.40%. An average of 25.4 million reads per sample were obtained for all tissue and salivary exosomes from OSCC patients as well as from the normal counterparts. Reads shorter than 15 nucleotides and with a quality score < 30 were excluded to ensure high quality of the sequencing results. Reads Per Million (RPM) was used to quantify miRNA expression level with false discovery rate (FDR) of ≤ 0.05, log2 fold change expression of ±2 and statistically significant abundance in each sample (p<0.05). We found miR-1307-5p to be significantly upregulated in tissue and salivary exosomal samples with a log2 fold change >2, and an average read count of 493 in OSCC salivary exosomal samples and 1297 in OSCC tissue samples. Interestingly, the control samples reported an average read count of 23 in salivary exosomal and 47 in tissue samples respectively. Given these low read counts in the controls, our results indicate that miR-1307-5p is significantly overexpressed in salivary exosomes and tumour tissues derived from OSCC patients as compared to their non-cancerous counterparts (Figure 2(a)). Thus, enhanced expression of miR-1307-5p in the primary tumour could be reflected in the salivary exosomal fractions, thus making it a potential liquid biopsy based biomarker for monitoring disease progression.

**Figure 2:**
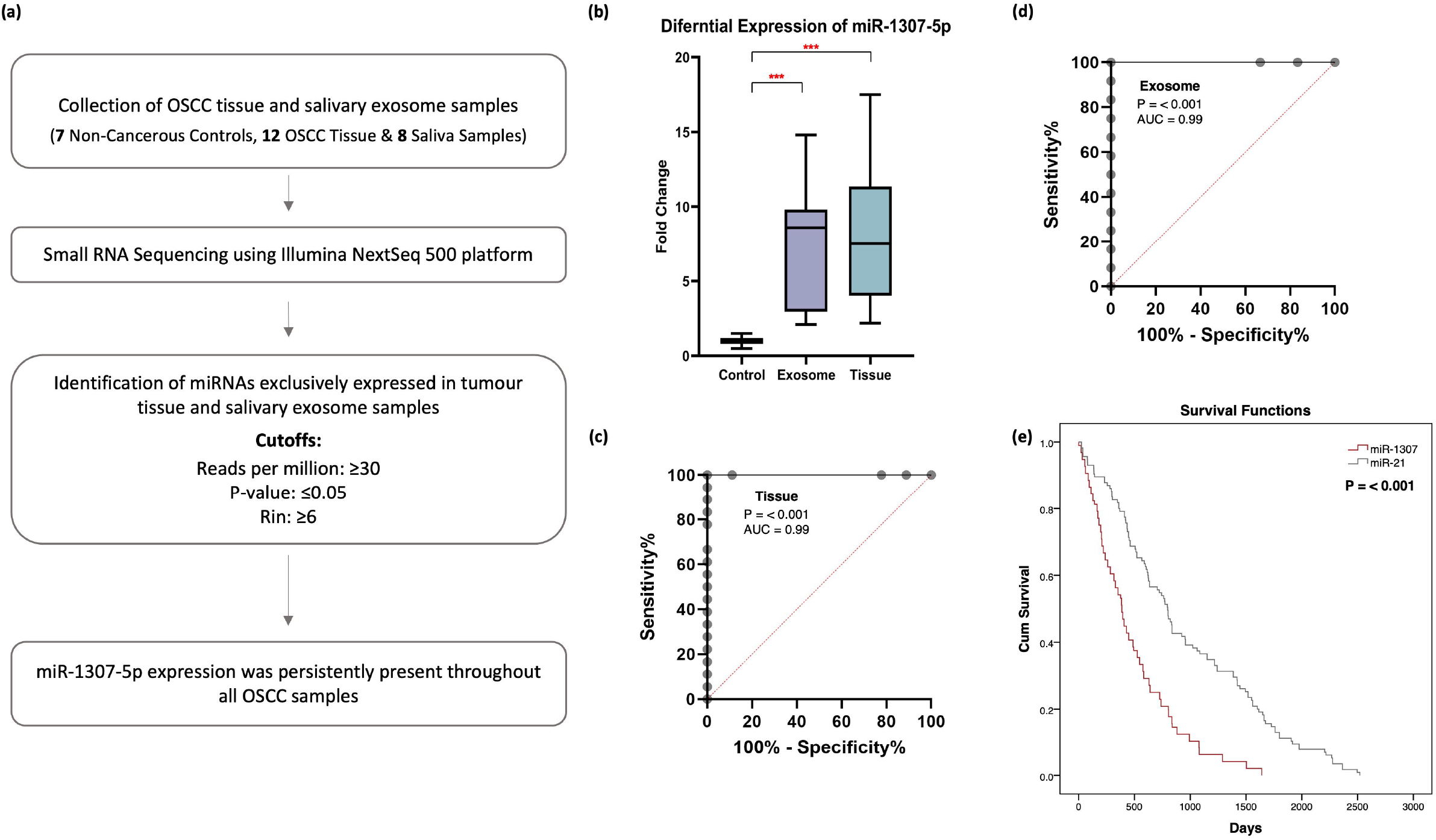
miR-1307-5p is differentially expressed and predicts poor overall survival in oral cancer patients. (a) Data analysis pipeline for small RNA sequencing. (b) Box plots representing the expression of miR-1307-5p in 15 salivary exosomal samples of OSCC patients and 19 OSCC tissue samples compared to healthy controls. The expression levels of miRNAs were estimated using real-time PCR. The combined measure of sensitivity and specificity miR-1307-5p was 99.99% across (c) tissue and (d) exosome samples (AUC:0.99, 95 % CI, p<0.001) Diagonal reference line acts as a performance measure of the diagnostic test. Note: AUC: Area Under the Curve, CI: Confidence Interval. (e) Kaplan–Meier plot of overall survival of miR-1307-5p and miR-21 generated from data available on TCGA_HNSC and data obtained from literature. The expression levels of miRNAs were estimated using real-time PCR. The data were normalized with U6 values, and the relative expression of miRNAs was analyzed using the ddCt method. Buccal scrapings and saliva samples obtained from healthy controls were used to calculate the relative expression of miR-1307-5p in OSCC tissue and salivary exosomes respectively. Error bars represent mean ± SD of three independent experiments (*p ≤ 0.05; **p ≤ 0.05; ***p ≤0.001).

### miR-1307-5p as a potential prognosticator in OSCC patients

Further we substantiated the expression profile of miR-1307-5p in the validation cohort using Real time PCR. An upregulation of miR-1307-5p was observed in OSCC tissue (FC:10.3±7.9, p-value: 0.02) and salivary exosomal samples (FC:7.3 ± 4.5, p-value: 0.0083) which was in agreement with the sequencing data (Figure 2(b)).

The predictive power of miR-1307-5p was estimated using logistic regression models, in which miRNA expression profile was used as predictor. Receiver operator characteristic (ROC) curves were determined and area under the curve (AUC) was considered. The ROC values for miR-1307-5p showed statistically significant diagnostic strength of this miRNA with sensitivity and specificity of 99.99% across all sample types (AUC:0.99, 95 % CI, p<0.001) (Figure 2(c,d)).

Several reports suggested that enhanced expression of miR-21 is associated with the poor overall survival rate in OSCC patients (13, 14). Hence to ascertain whether miR-1307-5p exhibited potential as a prognosticator, we conducted survival analysis of miR-21-5p with miR-1307-5p using their normalised expression value from TCGA datasets and literature survey (205 OSCC patients).

Our results suggested that miR-1307-5p could predict poor overall survival rate more effectively and significantly (50% patients demonstrated mortality within 17 months) compared to miR-21-5p (p-value<0.001) (Figure 2(e)). Collectively, these results suggest that miR-1307-5p indicates predictive power and patient outcome with high accuracy, and can efficiently function as a novel and independent prognostic biomarker for OSCC patients.

### miR-1307-5p demonstrates clinical association with disease progression, aggressiveness and therapeutic refractoriness

We further evaluated the relevance of enhanced miR-1307-5p expression with various clinicopathological parameters of OSCC patients. Increased levels of miR-1307-5p were observed in patients with high grade tumours (III/IV) in tissues (FC = 14.45 ± 9.76, p-value: 0.04) compared to low grade tumours (I/II) (FC = 5.37 ± 3.25). These findings were consistent in salivary exosomal samples of high grade (FC = 10.9 ± 5, p-value: 0.02) patients as compared to low grade OSCC patients (FC = 3.1 ± 1.2) (Figure 3(a)). Moreover, elevated levels of miR-1307-5p was observed in patients with locoregional aggressiveness (N1, N2, N3) in both tumour tissues (FC:17.5 ± 10, p-value: 0.03) and salivary exosomal samples (FC:13 ± 4, p-value: 0.0037) in comparison to those who did not report lymph node involvement (N0) (tissue FC = 7.5 ± 4.5; salivary exosome FC = 5 ± 2.7) (Figure 3(b)).

**Figure 3:**
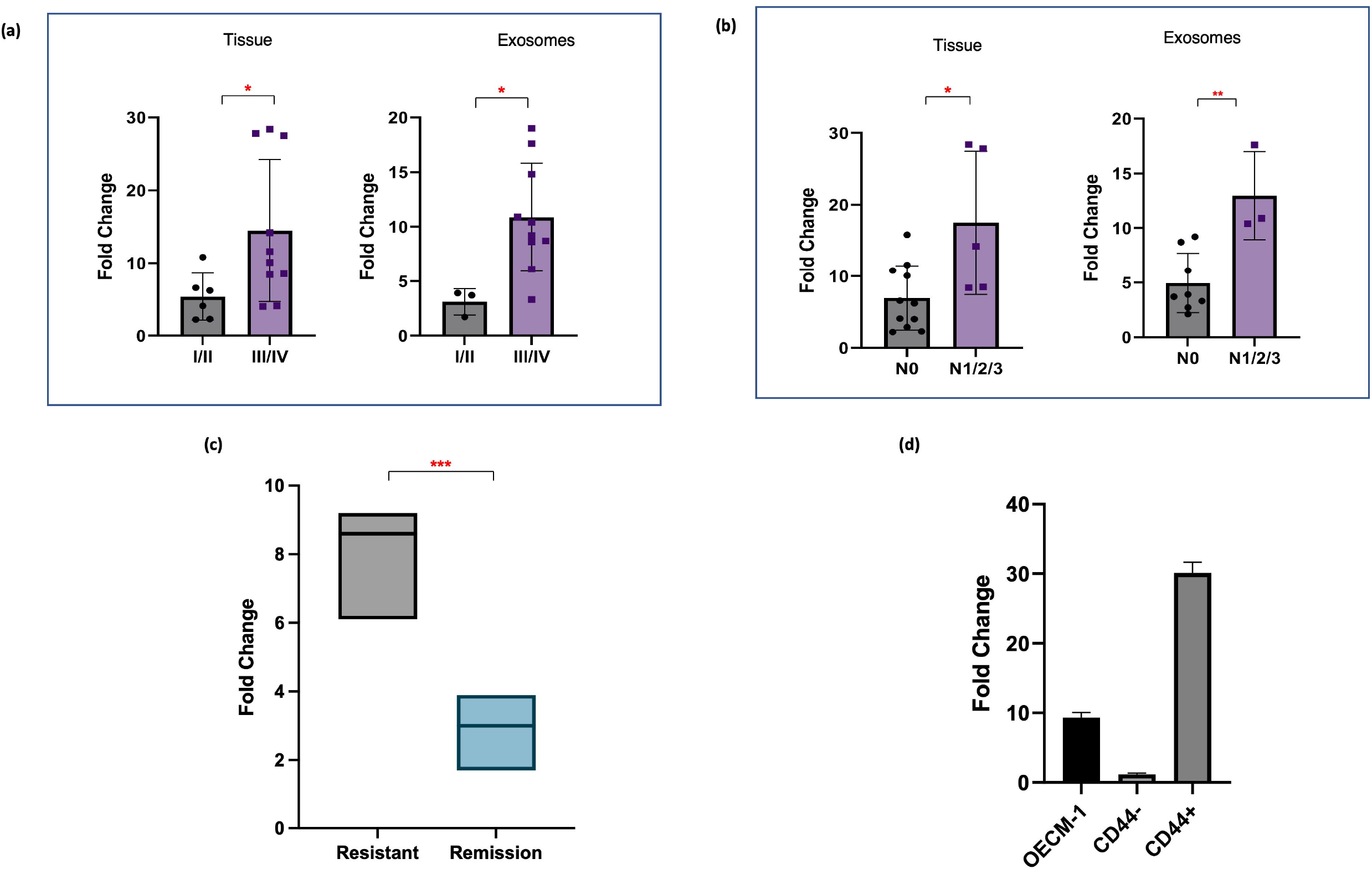
miR-1307-5p is overexpressed in late stage tumours and refractory, chemoresistant tumours. Representative box plots depict relative expression patterns of miR-1307-5p in (a) early stages (Stage I/II) vs late stages (Stage III/IV) of OSCC patient-derived tissue samples and salivary exosomal samples, (b) OSCC tissue and salivary exosome samples of patients with no nodal metastasis (N0) vs those with metastasis (N1/2/3), and (c) salivary exosomal samples of OSCC patients with recurrent tumours and those with complete remission. (d) Expression of miR-1307-5p in CD44+ and CD44-cells determined by qRT-PCR. The data were normalized with U6 values, and the relative miRNA levels were analyzed using the ddCt method. Buccal scrapings and saliva samples obtained from healthy controls were used to calculate the relative expression of miR-1307-5p in OSCC tissue and salivary exosomes respectively. Expression of miR-1307-5p in OECM-1, CD44-, and CD44+ samples was compared to buccal scrapings obtained from healthy controls. Error bars represent mean ± SD of three independent experiments (*p ≤ 0.05; **p ≤ 0.01; ***p ≤0.001).

Further, to assess the significance of miR-1307-5p on real time therapeutic monitoring, we analysed the expression levels of miR-1307-5p in chemoresistant OSCC patients compared to chemosensitive patients that showed complete disease remission post treatment. A significant increase in the expression levels of salivary exosomal miR-1307-5p was observed in the chemoresistant cohort (FC:4.82 ± 2.38, p-value:0.01) compared to patients with complete remission (FC:2.3 ± 1.2) (Figure 3(c)).Given that our analysis was conducted on a smaller patient cohort, further analysis on a larger cohort is warranted. Considering that cancer stem cells (CSCs) are a group of quiescent cells within the tumour bulk, that are believed to be responsible for tumorigenesis, resistance, relapse and poor clinical outcomes, we evaluated the expression of miR-1307-5p in CD44+ subpopulation derived from OECM1 cell line (15). CD44+ CSC subpopulation in OECM1 cells were isolated using immunomagnetic bead separation. miR-1307-5p expression was significantly upregulated in immuno-magnetically isolated CD44+ CSC subpopulation (FC: 31.3 ± 3.46,p=0.0001) as compared to CD44- (FC: 1.18 ± 0.16) fraction derived from OECM1 cell line (Figure 3(d)). Collectively, these findings clearly indicated that the presence of miR-1307-5p could be clinically associated with disease progression, local aggressiveness and chemotherapeutic refractoriness, thus making it an ideal prognosticator for OSCC patients. Thus, it became imperative to understand its miRNA-mRNA networks and underlying mechanism by which it regulates OSCC progression.

### Identification of differentially expressed genes in OSCC patients using RNA sequencing analysis

RNA-seq analysis of matched OSCC patients (that were utilised for small RNA sequencing) was conducted to identify mRNA targets and deciphering the molecular and functional mechanism of miR-1307-5p. Simultaneous profiling of miRNAs and mRNAs provides an opportunity to compare the gene expression of miRNAs and their target mRNAs without extra efforts in filtering out false positives from miRNA–target prediction. A total of 17.8-28.5 million reads were obtained from tissue samples. Further, an average of 62% of the reads mapped with the reference human genome (hg38) with high confidence. A total of 10660 differentially expressed mRNAs from tumour patient samples were identified as compared to their non-cancerous counterparts. On comparing these with the TCGA datasets, we found that a total of 5868 mRNAs were significantly expressed (log2FC= + 2; p<0.01) across these datasets irrespective of the varying clinico-pathological status of patients and discrepancy in sequencing techniques.

### Target gene prediction and functional analysis of miR-1307-5p

To identify the predicted target genes of miR-1307-5p TargetScan was used. Out of the predicted 120 targets, 43 genes were significantly expressed across the TCGA_HNSC dataset and the data generated from RNASeq in this study (Figure 4(a,b); Table 2). Since miRNAs are believed to negatively regulate their targets, 14 downregulated genes (Figure 4(c,d) were considered for further analysis (16, 17). A relative predicted KD cut-off of <-2 revealed 8 genes with the highest binding affinity to miR-1307-5p including *THOP1, PIM3, MGRN1, EHF, RNF4, ZNF726, GET4*, and *RNF114* with predicted relative KD values of −2.017, −3.493, −2.362, −3.068, −2.649, −2.107, −2.399, and −2.985 respectively (Figure 5(a)).

**Figure 4:**
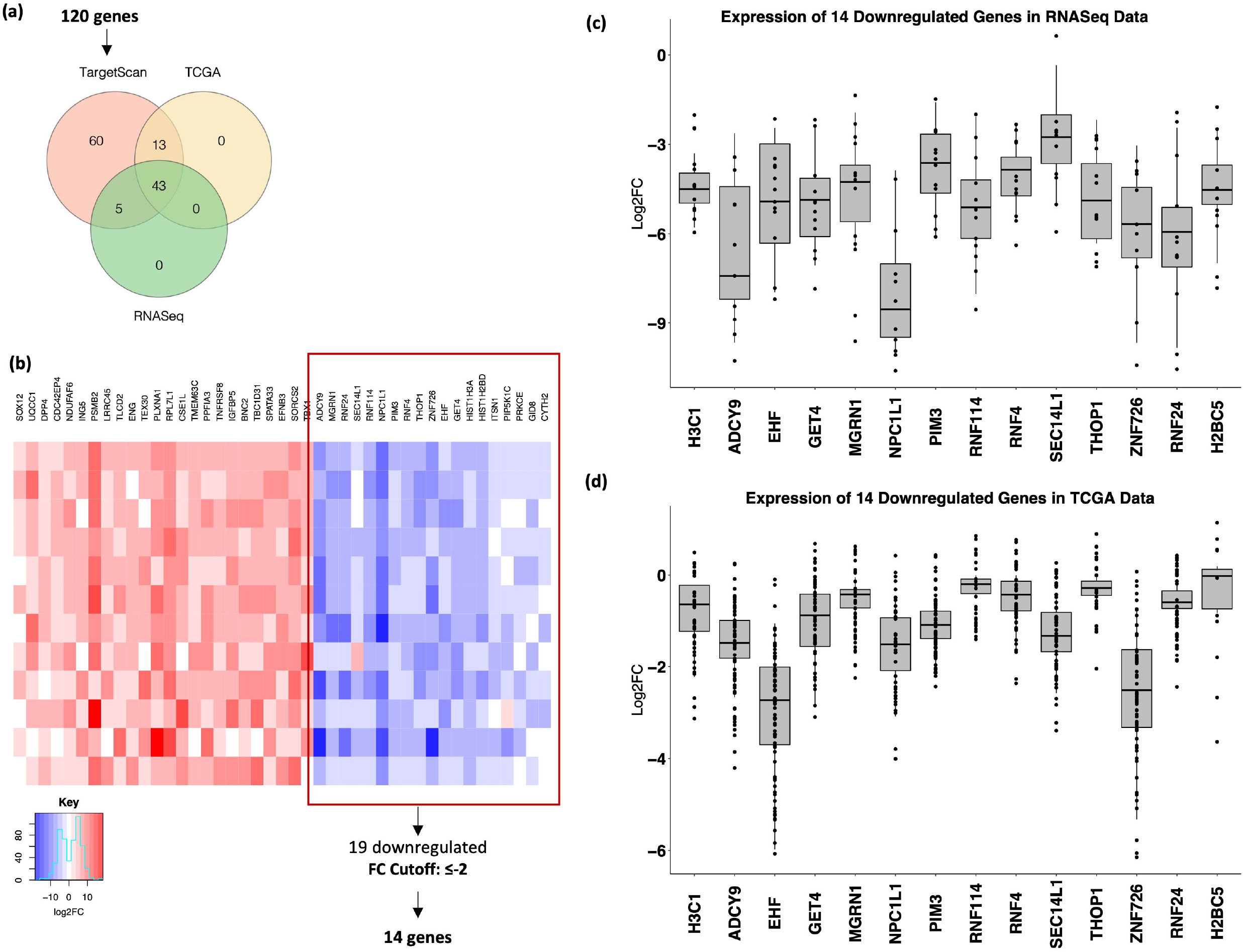
Validation of target genes of miR-1307-5p in OSCC. (a) Venn diagram depicting the selection of potential targets of mR-1307-5p via TargetScan 8, expression in TCGA_HNSC dataset and RNASeq expression generated in this study. 43 commonly expressed genes were identified out of which 30 were downregulated. A fold change cutoff of ≤ −2 revealed 14 significantly downregulated genes. (b) Heatmap of the commonly expressed 43 genes showed 30 downregulated genes out of which 14 genes passed the FC cutoff of ≤−2. Representative boxplots show the expression of the shortlisted *HIST1H3A (H3C1), ADCY9, EHF, GET4, MGRN1, NPC1L1, PIM3, RNF114, RNF4, SEC14L1, THOP1, ZNF726, RNF24*, and *HIST1H2BD (H2BC5)* in (c) RNASeq and (d) TCGA_HNSC datasets.

**Figure 5:**
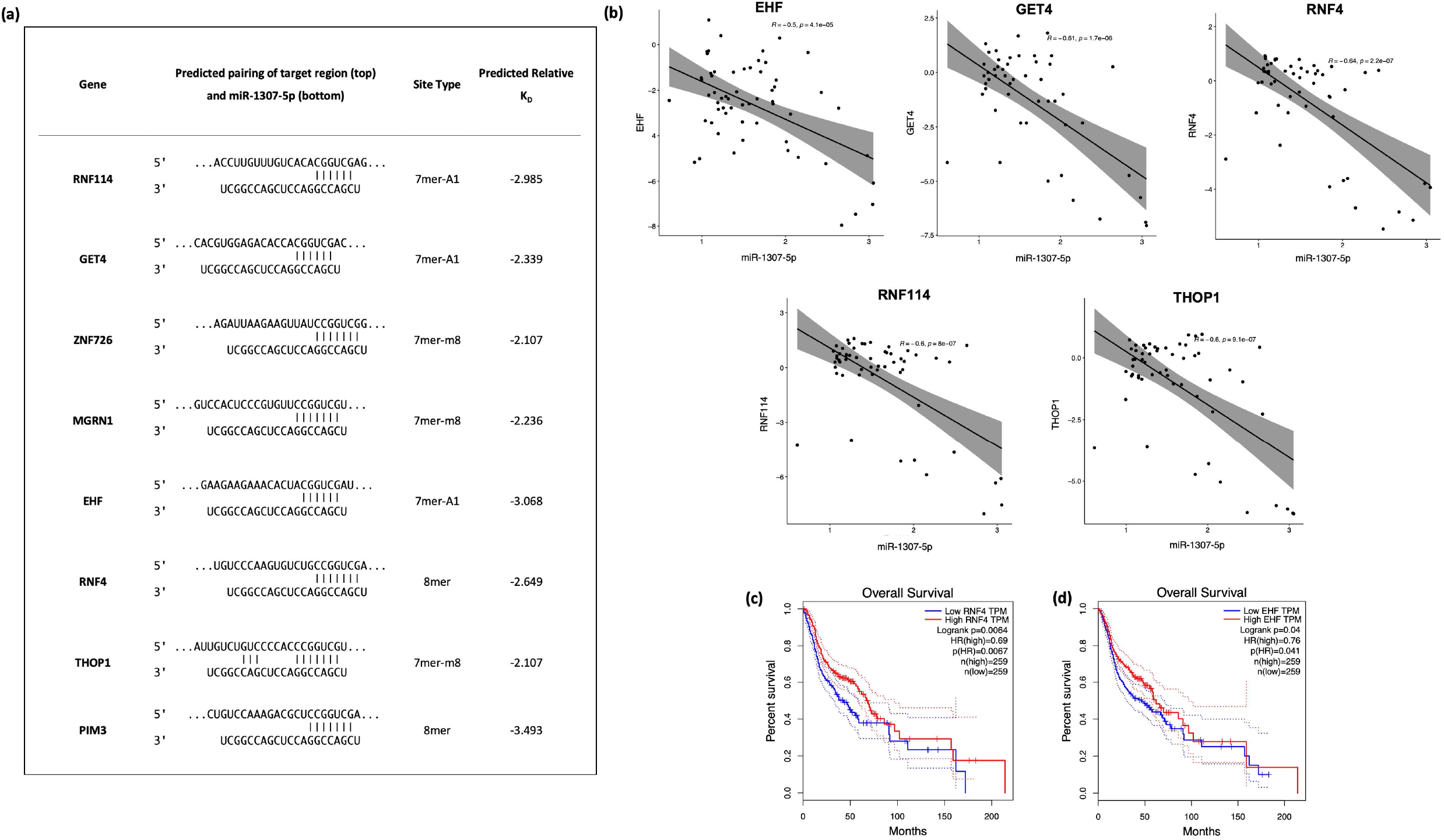
Predicted targets of miR-1307-5p show an inverse correlation with miR-1307-5p expression and predict overall survival in OSCC patients. (a) Table of predicted binding sites of target genes and miR-1307-5p along with their respective predicted KD values; (b) Pearson’s correlation analysis using RNASeq expression data and the expression data in TCGA_HNSC to identify inversely correlated genes. Overall survival analyses were performed using the GEPIA platform where patients with (c) RNF4 and (d) EHF expression above the median are indicated by red lines, and patients with gene expression below the median are indicated by black lines. Log-rank p < 0.05 was considered to indicate a statistically significant difference.

**Table 2:** Expression of the predicted target genes of miR-1307-5p in TCGA dataset and the RNASeq data generated in this study.

| Gene | LOG2FC |  |
| --- | --- | --- |
|  | TCGA | RNASeq |
| **Downregulated** |  |  |
| *NPC1L1* | -6.24 | -7.32 |
| *ZNF726* | -3.05 | -5.67 |
| *HIST1H3A* | -4.89 | -4.65 |
| *ADCY9* | -4.23 | -3.98 |
| *EHF* | -2.51 | -3.06 |
| *RNF4* | -2.52 | -2.89 |
| *HIST1H2BD* | -2.95 | -2.74 |
| *RNF24* | -2.67 | -2.45 |
| *GET4* | -2.41 | -2.41 |
| *PIM3* | -2.77 | -2.34 |
| *GID8* | -1.68 | -2.27 |
| *THOP1* | -2.27 | -2.19 |
| *RNF114* | -2.65 | -2.02 |
| *MGRN1* | -2.29 | -1.95 |
| *SEC14L1* | -2.34 | -1.82 |
| *CYTH2* | -1.18 | -1.72 |
| *PIP5K1C* | -1.98 | -1.50 |
| *ITSN1* | -2.36 | -1.29 |
| *PRKCE* | -1.93 | -0.88 |
| **Upregulated** |  |  |
| *SOX12* | 1.56 | 2.19 |
| *UQCC1* | 2.17 | 2.46 |
| *ENG* | 3.54 | 2.79 |
| *NDUFAF6* | 2.95 | 3.30 |
| *TLCD2* | 3.50 | 3.31 |
| *ING5* | 3.03 | 3.37 |
| *LRRC45* | 3.22 | 3.40 |
| *DPP4* | 2.60 | 3.63 |
| *CDC42EP4* | 2.79 | 3.63 |
| *PSMB2* | 3.19 | 3.90 |
| *CSE1L* | 3.94 | 3.97 |
| *TEX30* | 3.56 | 4.05 |
| *RPL7L1* | 3.86 | 4.06 |
| *TMEM63C* | 4.18 | 4.11 |
| *PPFIA3* | 4.83 | 4.72 |
| *PLXNA1* | 3.86 | 4.73 |
| *IGFBP5* | 7.18 | 5.33 |
| *TNFRSF8* | 6.07 | 6.78 |
| *SPATA33* | 7.92 | 8.27 |
| *EFNB3* | 8.48 | 8.50 |
| *BNC2* | 7.29 | 8.56 |
| *TBC1D31* | 7.77 | 8.98 |
| *SORCS2* | 8.78 | 9.54 |

miRNAs regulation of gene expression could result in an inverse correlation between miRNA and its target (18). To measure the inverse correlation between miR-1307-5p expression and the expression of the 8 target genes, we performed Pearson correlation using RNA seq and TCGA HNSC expression data. Out of the identified 8 genes, *THOP1* (p=9.1e-0.7)*, EHF* (p=4.1e-0.5)*, RNF4* (p=2.2e-0.7)*, GET4* (p=1.7e-0.6), and *RNF114* (p=8e-0.8) showed significant correlation with the expression of miR-1307-5p *(p<0.0001)* (Figure 5(b)). Amongst these genes, lower expression RNF4 (p= 0.0064) (Figure 5(c)) and EHF (p= 0.04) (Figure 5(d)) correlate with poor overall survival in HNSCC patients in TCGA datasets.

## Discussion

Exosomes have been intensively studied for disease progression, aggressiveness and therapeutic monitoring in OSCC (19). Moreover, the presence of exosomes in biological fluids such as saliva, cerebrospinal fluid, urine, and blood is providing a new source for biomarkers for OSCC (20, 21). However, until now no standard method has been established for isolation and characterisation of exosomes. Various surface markers such as CD9, CD63, CD81 and Alix have been identified as membrane protein markers for exosomes. Many reports have suggested that CD47 is also present on exosomes; however, whether CD47 plays a direct or an indirect role in determining exosomes remains unclear (22). Our study shows a significant CD47 expression in salivary exosomes of OSCC patients, demonstrating a great potential of individual exosome profiles in biomarker discovery. This is the first study to show expression of CD47 on salivary exosomes and further research is required to understand the role of CD47 as a probable surface marker for exosomes.

The purpose of this study is to identify an exclusive biomarker for OSCC prognosis using salivary exosomal miRNAs. We found enhanced expression of miR-1307-5p in tumour tissues and exosomes as an independent risk predictor and prognosticator for OSCC patients. Concurring expression profiles of miR-1307-5p in salivary exosomes and tissue samples clearly suggests the crucial role and potential clinical utility of this non-invasive prognostic biomarker using liquid biopsy approach. This miRNA demonstrated a significant clinical association with disease progression, locoregional aggressiveness and chemotherapeutic refractoriness. Abundance of miR-1307-5p in CD44+ stem cell subpopulation derived from OECM-1 cell line substantiated our clinical findings and were indicative of the fact that identified miRNA has a role in regulating the self-renewal and maintenance of CD44+ subpopulation that is responsible for incurring disease aggressiveness and therapeutic resistance in OSCC patients. Few studies have reported the role of miR-1307-5p as an important prognosticator and risk predictor in various malignancies specifically using non-invasive approaches. Xinyue Du (2020) reported the role of miR-1307-5p in proliferation and invasion by targeting TRAF3 and activating NF-κB/MAPK pathway in lung adenocarcinoma (23). Thus, identifying miR-1307-5p/TRAF3/NF-κB/MAPK axis as a potential prognostic and therapeutic target. miR-1307 was found to be upregulated in chemoresistant ovarian tumour tissues and cell lines compared to chemosensitive population, suggesting the role of miR-1307 in development of chemoresistance in ovarian cancer (24). Moreover, high levels of miR-1307 in serum exosomes of ovarian cancer patients showed independent diagnostic power and were associated with tumour staging (25). These results support our findings of the role of miR-1307-5p in disease progression, locoregional aggressiveness and chemoresistance. However, the role of miR-1307-5p remains completely unexplored in head and neck/oral cancers. To the best of our knowledge this is the first study reporting the prognostic potential of miR-1307-5p in OSCC using liquid biopsies.

Given that, miRNAs regulate target mRNAs by binding to the 3’ UTR of target genes in a post-transcriptional manner, establishing the interrelationship of miR-1307-5p and its target genes would enhance our understanding of the molecular mechanism and provide potential therapeutic targets for oral cancers (26). We identified 43 significantly expressed putative target genes of miR-1307-5p using transcriptome sequencing analysis, TCGA datasets and TargetScan software. Several studies have reported that miRNAs result in suppressing the gene expression, by conducting mRNA degradation and by inhibiting protein translation or degrading the polypeptides through binding complementarity to 3’ UTR of the target mRNAs. Hence, in this study we focused on the 14 downregulated target genes. Amongst these, 5 genes *(THOP1, EHF, RNF4, GET4, RNF114)* were found to have a significant association and formed strong miRNA-mRNA networks with miR-1307-5p based on significant expression patterns, alignment scores with the identified miRNA and Pearson’s correlation scores using miRNA-mRNA expression patterns.

EHF is a transcription factor that has a vital role in cell differentiation, and tumour-initiating and metastatic capability by conferring a CSC-like phenotype (27). In HNSCC, loss of EHF either by point mutations or altered expression patterns is reported to promote aggressiveness of the disease by inducing EMT phenomena or by targeting regulators of redox homeostasis such as NRF2 and SOX2 (28, 29). These reports are in concordance with our findings and suggest a crucial role of EHF in promoting progression and aggressiveness in OSCC tumours.

In this study, out of the 5 top genes targeted by miR-1307-5p, two genes belong to the E3 ubiquitin ligase family. Deregulated E3 ligases have been reported to confer uncontrolled proliferation leading to malignant transformation, progression and therapy resistance (30). RNF4 is associated with the double strand break repair mechanism and is responsible for inducing degradation of altered target proteins via proteasomes. RNF4 has been reported to promote ubiquitination of PML bodies which leads to proteasome degradation (31). Moreover, deregulation of PML bodies leads to activation of PPARδ mediated fatty acid oxidation pathways, which help in cancer stem cell maintenance, leading to disease progression and therapeutic refractoriness (32, 33). Another ligase, RNF114 has been reported to promote ubiquitination and degradation of p21 (34). p21 is an important regulator of the cell cycle. Stabilisation of p21 leads to cell cycle arrest and ultimately promotes apoptosis. This study demonstrated a significant downregulation of RNF4 and RNF114 in OSCC patients by miR-1307-5p. Suppression of these genes would not only impede apoptosis via cell cycle deregulation but also would promote cancer stem cell maintenance thereby leading to disease aggressiveness and therapeutic refractoriness.

THOP1 is a characteristic metallopeptidase with a HEXXH zinc binding motif and associated with metabolism of several neuropeptides’ acids, such as bradykinin, gonadotropin releasing hormone, opioids and neurotensin (35). Hydrolysis of Bradykinin by THOP1 induces accumulation of intracellular calcium ions (Ca2+), thus leading to Ca2+ homeostasis which incur tumour initiation, angiogenesis, progression and metastasis (36). Thus, further research on degradation of neuropeptides by THOP1 will lead to interesting models for future investigations in the development of cancer. Lastly, GET4 is one of the factors of the BCL2-associated athanogene 6 (BAG6) chaperone complex and it functions as a regulator for the nucleo-cytoplasmic transport of BAG6 (37). In one of the recent studies by Koike K et al., GET4 was identified as a novel colorectal cancer driver gene that promoted tumor growth by facilitating cell cycle progression (38). Collectively, these mRNA regulated by miR1307-5p modulate vital biological and functional processes such as cell cycle, proliferation, angiogenesis, EMT and CSC self-renewal (Figure 6). Interestingly, to the best of our knowledge, apart from EHF, there are no studies reporting the association of *THOP1, RNF4, GET4* and *RNF114* with oral cancers.

**Figure 6:**
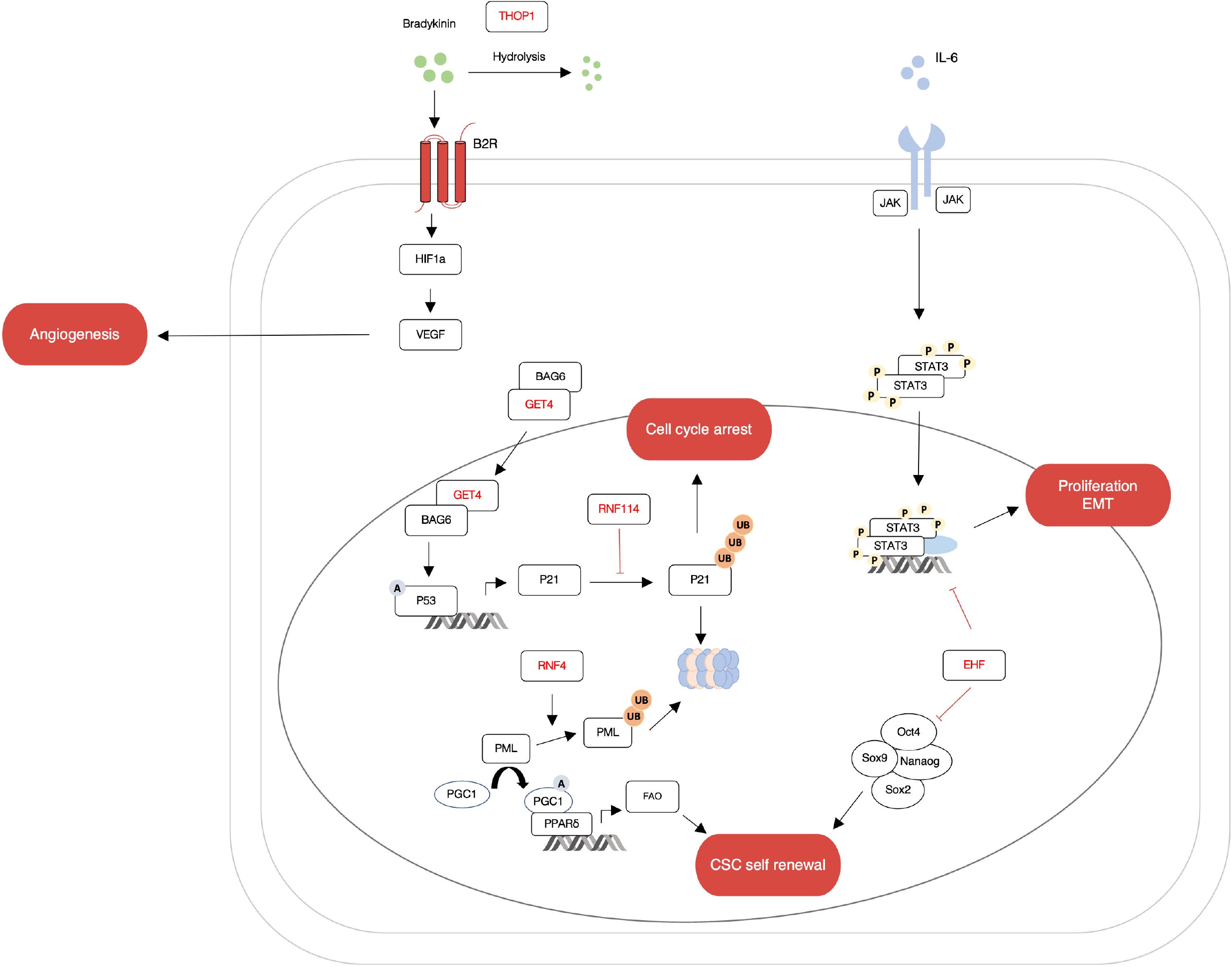
miR-1307 regulates important genes involved in OSCC progression. The figure shows a schematic representation of the relationships between putative genes targeted by miR-1307. Many of these genes modulate important cellular processes like cell proliferation, apoptosis, angiogenesis and maintenance of cancer stem cells.

This study is a preliminary attempt to identify the role of miR-1307-5p as a potential prognosticator in OSCC using liquid biopsy techniques and to elucidate the plausible mechanism by which this miRNA regulates mRNA targets (*THOP1, EHF, RNF4, GET4, RNF114*) thereby inducing disease aggressiveness and therapeutic refractoriness in OSCC patients. A better understanding of the underlying mechanisms and functional role of these identified genes via miR-1307-5p will help us identify novel prognostic markers and therapeutic targets for oral cancers.

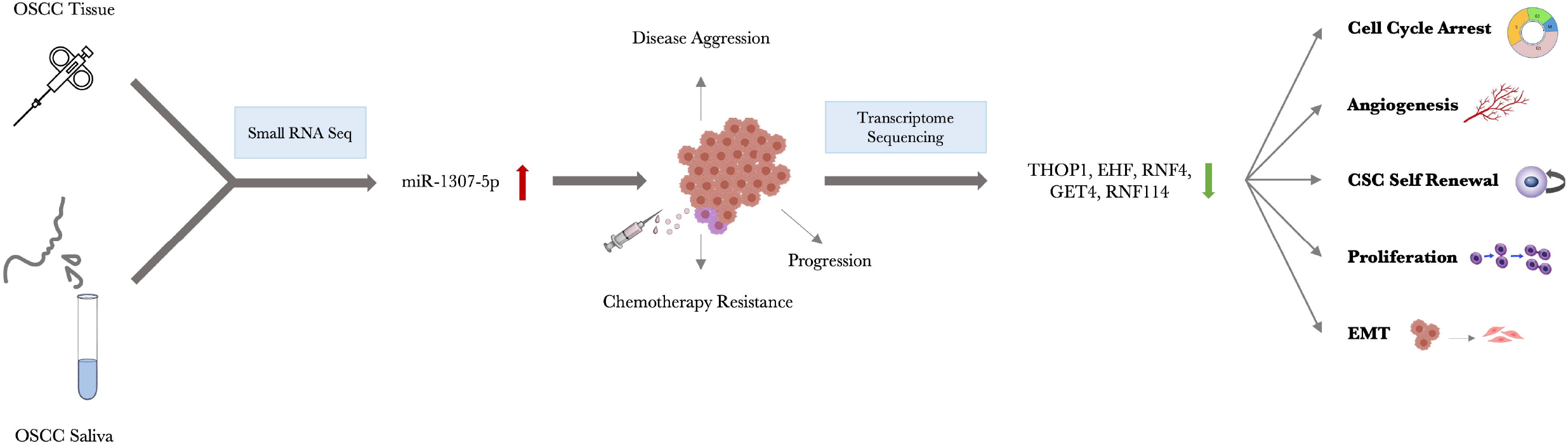

## Acknowledgements

The authors are grateful to Prof. Rakesh Rawal, Gujarat University for kindly providing access to the NanoSight LM10. We acknowledge, Dept. of Bioinformatics, Gujarat University for providing computational resources for data analysis.This facility was funded by the Gujarat State Biotechnology Mission and Gujarat Council on Science and Technology, Government of Gujarat. We appreciate the kind support from Beckman Coulter Inc for providing us with CytoFLEX-LX flow cytometer and its consumables on a trial basis.

## Ethics approval statement

All procedures of human samples were conducted after the approval of the HCG Cancer Centre’s Ethics Committee (ECR/92/Inst/GJ/2013/RR-16) for human subject research. This study also complied with the guidelines set forth by the Declaration of Helsinki (2008). All patients provided written informed consent for their participation in the study and their identities have been anonymised.

## Author contributions

KP and DM: provided clinical samples and clinical assistance; SP, DM, KP, and VT: conceived and designed the experiments; AP and PP: conducted the experiments; AP and SP: performed bioinformatic analysis; AP, SP and PP: performed data analysis; AP, SP and VT: wrote the manuscript; SP and VT: edited the manuscript.

